# Structure-derived synthetic sequences guide a protein language model toward metalloproteins

**DOI:** 10.64898/2026.04.30.722007

**Authors:** Giulia Peteani, Gianmattia Sgueglia, Thomas Lemmin, Marco Chino

## Abstract

**Motivation:** Protein language models (pLMs) capture evolutionary sequence constraints but are limited in modeling underrepresented functional classes due to training data imbalance. Metalloproteins constitute a fundamental but sparsely represented class in sequence databases. We therefore assess whether structure-conditioned synthetic sequences can be used to specialize pLMs toward metal-binding functionality.

**Results:** We fine-tuned the generalist model ProtGPT2 on synthetic sequences generated by the inverse-folding model ProteinMPNN, constructing training sets with controlled variation in size and diversity. Fine-tuning increased recovery of canonical metal-binding motifs from 43% in the baseline model to 91% in the fine-tuned models. Generated sequences retained high predicted structural confidence and structural similarity to known folds, despite low sequence identity. Analysis of latent representations from ProtGPT2 indicated that fine-tuned models occupy distinct regions of embedding space relative to both the baseline model and structure-conditioned sequences, consistent with partial incorporation of structural constraints while preserving sequence diversity. A multi-step filtering pipeline applied to sequences lacking canonical motifs identified candidate metal-binding sites in four-helical bundle topologies not detected in a non-redundant subset of Protein Data Bank structures or in AlphaFold-predicted proteomes.

**Availability and implementation:** Code, trained models, and datasets are available at: https://doi.org/10.5281/zenodo.18672158 and https://huggingface.co/gsgueglia.

## 1. Introduction

Protein language models (pLMs), including ProtGPT2^1^, ProGen2^2^, and ESM-2^3^, capture evolutionary and biophysical constraints of amino acid sequences by learning statistical patterns from large sequence databases^4,5^. These models can generate realistic sequences^1–3,6^, predict structures^3,7–9^ and stability^10^, and reflect residue-level dependencies relevant to function and evolution^5^. However, their ability to explore specialized protein families is limited^11,12^, and fine-tuning is generally the main strategy to overcome such limitations^13^. For many functionally annotated families, including metalloenzymes, ligand-binding domains, and membrane receptors, the number of experimentally characterized sequences is small, restricting access to high-value regions of sequence space.

Synthetic data can mitigate dataset limitations and enable model specialization^14–20^. In protein modeling, however, generating informative sequences is challenging because they must satisfy folding constraints while preserving functional features. One approach is to generate sequences from known structures using inverse-folding models^21,22^ which condition sequence design on a fixed backbone and encode structural constraints in sequence space. Sampling diverse sequences from selected templates can thus produce datasets that retain structural motifs and functional sites while expanding beyond the variation observed in natural databases.

Metalloproteins provide a stringent test case for this strategy. Approximately one third of proteins bind metal ions, supporting catalysis, electron transfer, sensing, and regulation across all domains of life^23–26^. Since metal binding depends on both coordination motifs and structural context, metalloproteins tightly couple sequence, structure, and function ^27–29^. Despite their prevalence, they remain challenging to model due to uneven annotation and the difficulty of distinguishing functional metal-binding sites from incidental coordination ^25,26,30,31^. This limitation constrains deep learning approaches that rely on large, accurately labeled datasets.

Several deep learning methods have been developed for metalloproteins, including metal-binding site detection^32–41^, coordination geometry prediction^41,42^, and metal classification^35,38,43–45^. However, these approaches are primarily predictive and do not address the generation of novel metalloprotein sequences.

In this study, we evaluate whether structure-derived synthetic sequences can be used to specialize protein language models (pLMs) toward functionally constrained regions of sequence space. We fine-tune the foundation model ProtGPT2 on synthetic proteins generated by ProteinMPNN, using metalloproteins as a biologically relevant test case. By comparing fine-tuned models to the base ProtGPT2 across sequence properties, predicted structural confidence, and metal-binding features, we assess how synthetic fine-tuning alters the generative distribution. Our results show that this approach enriches metal-binding–related features while preserving sequence diversity and generating sequences with low similarity to known proteins, while also exposing limitations that highlight the need for improved fine-tuning strategies and more accurate functional supervision.

## 2. Results

### Structure-derived synthetic data for model specialization

We fine-tuned ProtGPT2 using synthetic metalloprotein sequences generated with ProteinMPNN^21^ and evaluated the resulting models on de novo sequence generation. Metalloproteins provide a convenient experimental setting due to their well-defined coordination geometries and the availability of quantitative evaluation metrics, including metal-binding site recovery and structural fold consistency.

To examine how dataset size and chemical diversity affect specialization, ProtGPT2 was fine-tuned on three synthetic datasets that differ in metal-binding complexity and sequence volume (Figure 1). The α dataset focused on transition-metal–binding domains (Co, Ni, Fe, Mn, Cu, Zn, Cd) and was generated from 370 structural scaffolds, yielding approximately 70,000 synthetic sequences. The β and γ datasets expanded the chemical space to include alkaline earth metals (Ca, Mg) and were generated from approximately 2,000 scaffolds. The β dataset comprised approximately 140,000 sequences, whereas the γ dataset scaled to approximately 1.4 million sequences. In all cases, ProteinMPNN performed fixed-backbone design, preserving the native metal-coordinating residues.

**Figure 1.**
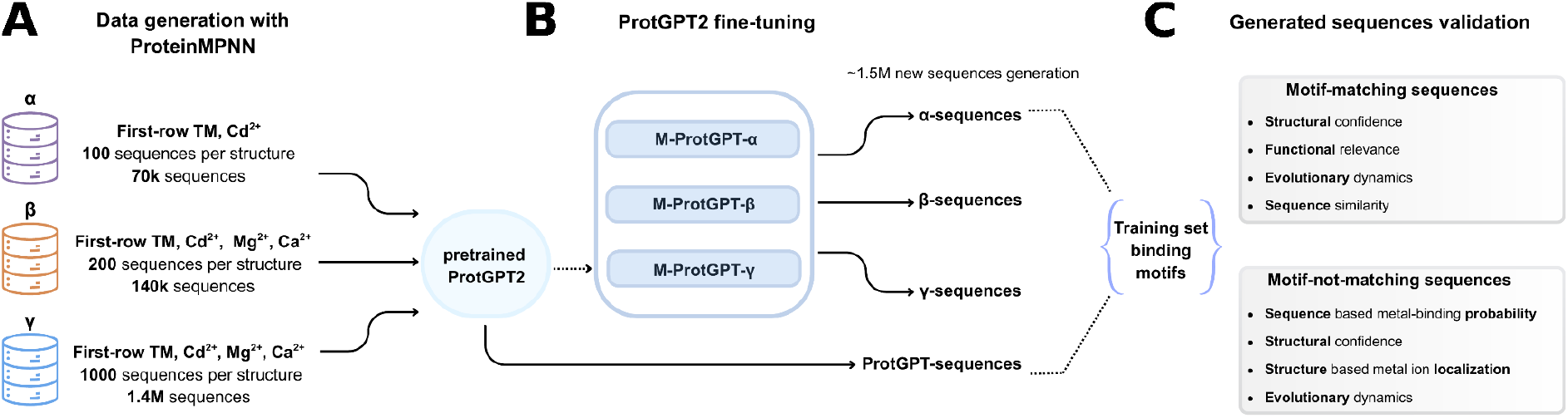
Workflow for structure-derived synthetic data fine-tuning of ProtGPT2. (A) ProteinMPNN was used to generate structure-conditioned sequences for three datasets (α, β, γ) differing in size and metal-binding site composition. (B) The pretrained ProtGPT2 model was fine-tuned on each dataset, yielding three specialized variants (M-ProtGPT2-α, -β, -γ), which were used to generate de novo protein sequences. (C) Generated sequences were stratified based on the presence of known metal-binding motifs and evaluated using sequence-based and structure-based metrics, including structural confidence, metal-binding site features, sequence similarity, and evolutionary properties.

Each dataset was used to fine-tune the pretrained ProtGPT2 model, producing the M-ProtGPT2-α, M-ProtGPT2-β, and M-ProtGPT2-γ variants. For evaluation, we generated approximately 1.3, 1.6, and 1.4 million de novo sequences from the α, β, and γ models, respectively, alongside 1.7 million sequences from the base ProtGPT2 model. All sequences were generated at a target length of approximately 100 amino acids and used as a common test set for subsequent analyses.

As expected, analysis of sequence diversity shows that ProtGPT2 generates broadly diverse sequences (Supplementary Figure S1). In contrast, fine-tuned models, especially M-ProtGPT2-β and -γ, produced more internally similar outputs, consistent with a focused exploration of sequence space shaped by metal-binding and structural constraints rather than a loss of generative capacity.

### Binding motif preservation and sequence quality

To quantify the impact of fine-tuning on metal-binding sequence features, all generated sequences were screened for known metal-binding motifs using regular expressions derived from canonical motifs in the training datasets. Fine-tuning markedly increased the proportion of sequences containing these motifs, with motif matches observed in 77% (1.1 M) of M-ProtGPT2-α sequences, 91% (1.3 M) of M-ProtGPT2-β sequences, and 85% (1.2 M) of M-ProtGPT2-γ sequences, compared to 43% (0.64 M) for the baseline ProtGPT2 model.

We next assessed the structural quality of motif-containing sequences, focusing on M-ProtGPT2-γ since it was trained on the largest and most chemically diverse dataset. Motif-matching sequences for representative metalloprotein prototypes were folded with ESMFold^3^ (Supplementary Table S1), and predicted pLDDT scores were compared to those of motif-matching sequences generated by ProtGPT2. Across all metalloprotein classes, the pLDDT distribution of M-ProtGPT2-γ sequences shows a modest but consistent shift toward higher values relative to the baseline (Figure 2A) Predicted structures of motif-matching M-ProtGPT2-γ sequences were also compared with their reference PDB structures using Foldseek^46^ (Supplementary Figure S2). Across multiple metalloprotein classes, γ-derived sequences more frequently adopted folds similar to the reference structures than sequences generated by the baseline model (Figure 2B), especially for hydrolases, oxidases, zinc finger proteins, and blue copper proteins.

**Figure 2.**
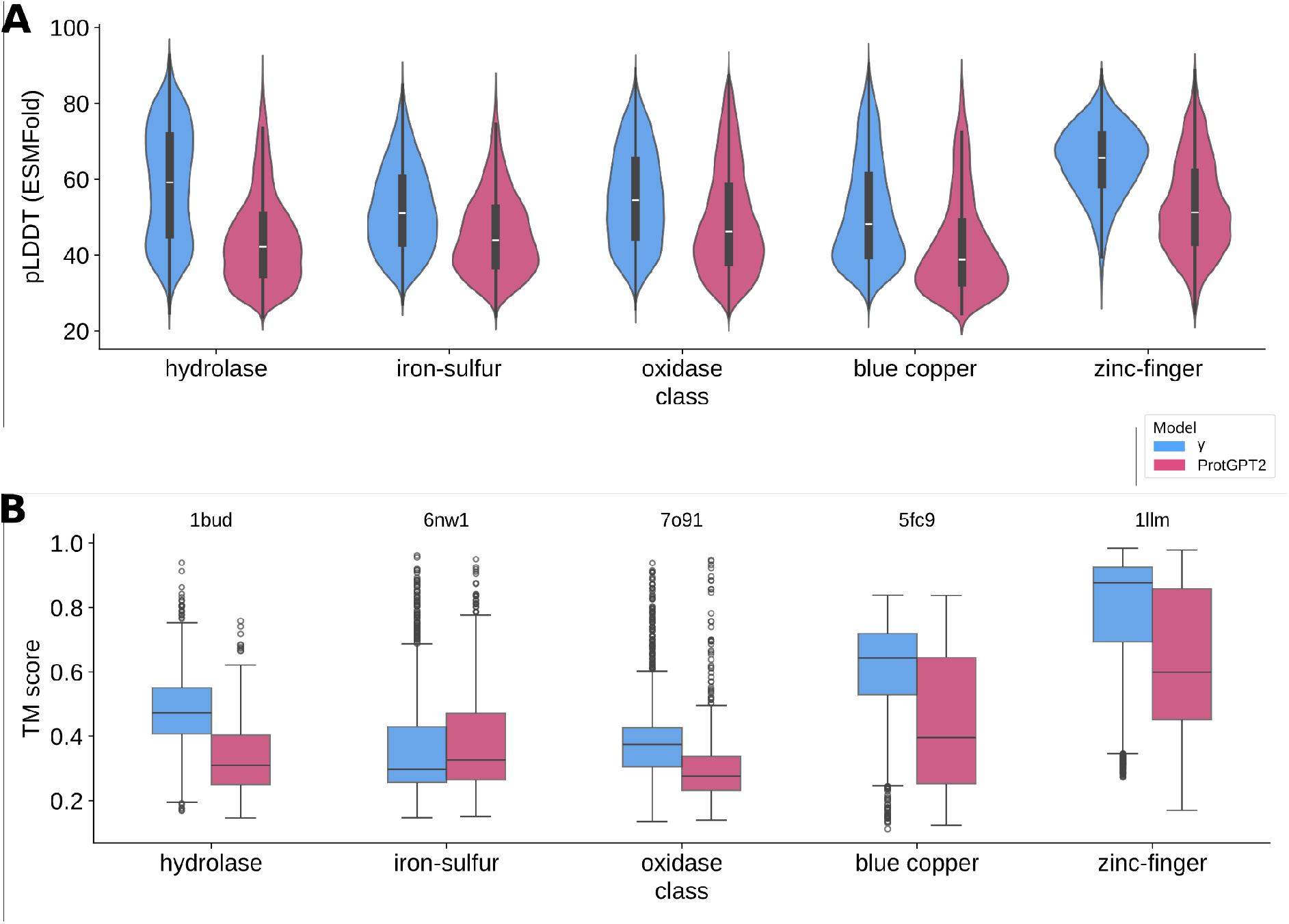
Structural evaluation of metal-binding motif–matching sequences generated by ProtGPT2 and M-ProtGPT2-γ. (A) Violin plots showing the distribution of ESMFold-predicted pLDDT scores for metal-binding motif–matching sequences generated by the M-ProtGPT2-γ and baseline ProtGPT2 models across five representative metalloprotein classes. (B) Box plots showing the distribution of TM-scores comparing predicted structures of motif-matching sequences to the corresponding reference PDB structures for each class. Results are shown for M-ProtGPT2-γ and ProtGPT2. The reference PDB ID used for each class is indicated above the corresponding boxplots.

In the zinc finger, hydrolase, and blue copper classes, sequences with high structural similarity (TM-score > 0.8) exhibited low sequence identity (< 0.3), indicating that the model can generate structurally similar yet sequence-divergent variants (Figure 3). In contrast, iron–sulfur and oxidase classes showed lower structural recovery, with most generated sequences falling below a TM-score of 0.5 irrespective of sequence identity. Notably, these classes were also represented by substantially fewer motif-matching sequences, limiting the statistical power of the analysis and suggesting reduced coverage for these coordination chemistries.

**Figure 3.**
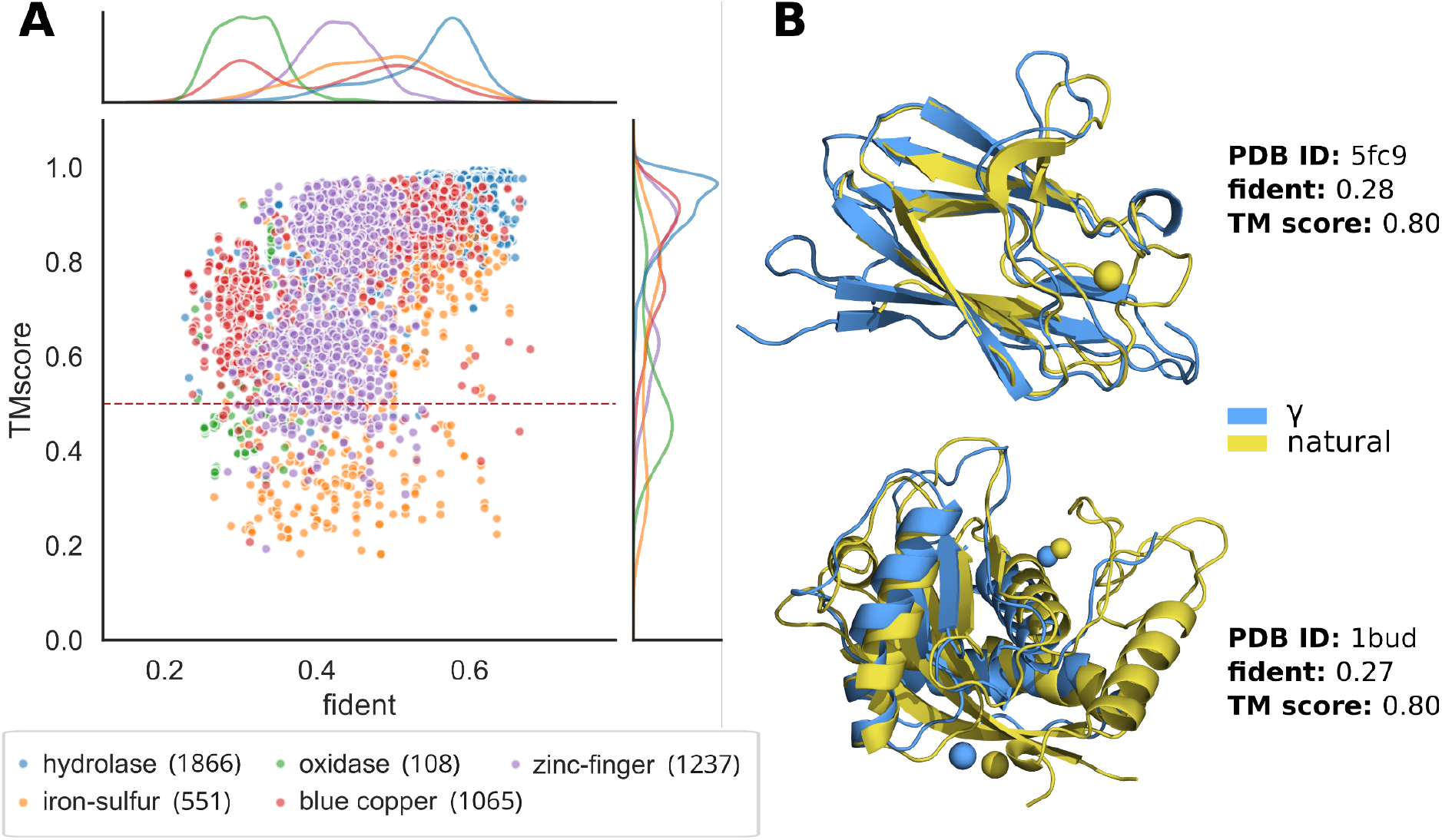
Structural similarity of metal-binding motif–matching γ-sequences to reference proteins. (A) Relationship between Foldseek TM-score and MMseqs sequence identity (fident) for motif-matching M-ProtGPT2-γ sequences relative to their corresponding reference PDB structures across five metalloprotein classes. The number of motif-matching sequences for each reference structure is reported in parentheses. The red dashed line indicates TM = 0.5, a commonly used threshold for fold-level structural similarity. (B) Representative structural alignments illustrating high structural similarity despite low sequence identity. Experimentally determined reference structures are shown in yellow (1bud, hydrolase; 5fc9, blue copper), and corresponding generated γ-sequences folded with Chai-1 are shown in blue, with coordinated zinc ions displayed. Both pairs exhibit TM-scores of 0.80 with sequence identities (fident) of 0.27 and 0.28, respectively.

To examine how generated sequences relate to natural, redesigned, and baseline ProtGPT2 sequences in the embedding space, we employed Evolocity^47^, which infers directional relationships between sequences from local embedding geometry and similarity patterns. Analyses were restricted to zinc-finger sequences with TM-scores > 0.5 relative to the reference PDB, focusing on structurally plausible variants while maintaining sufficient sequence diversity. Zinc-fingers represents a well-defined fold-function category and was the only metal-binding protein for which baseline ProtGPT2 produced sufficient sequences to enable Evolocity analysis.

ESM-1b embeddings projected onto the first two principal components separated ProtGPT2 and M-ProtGPT2-γ sequences, while synthetic redesigns partially overlapped the edge of the γ cluster (Figure 4A). Evolocity inferred two main trajectories from baseline ProtGPT2: one toward γ-generated and redesigned sequences, and one toward natural sequences. This bifurcation suggests that the model assigns higher local likelihood to both γ-generated and natural sequences relative to baseline ProtGPT2 outputs, while distinguishing between redesign-driven and naturally evolved sequence solutions.

**Figure 4.**
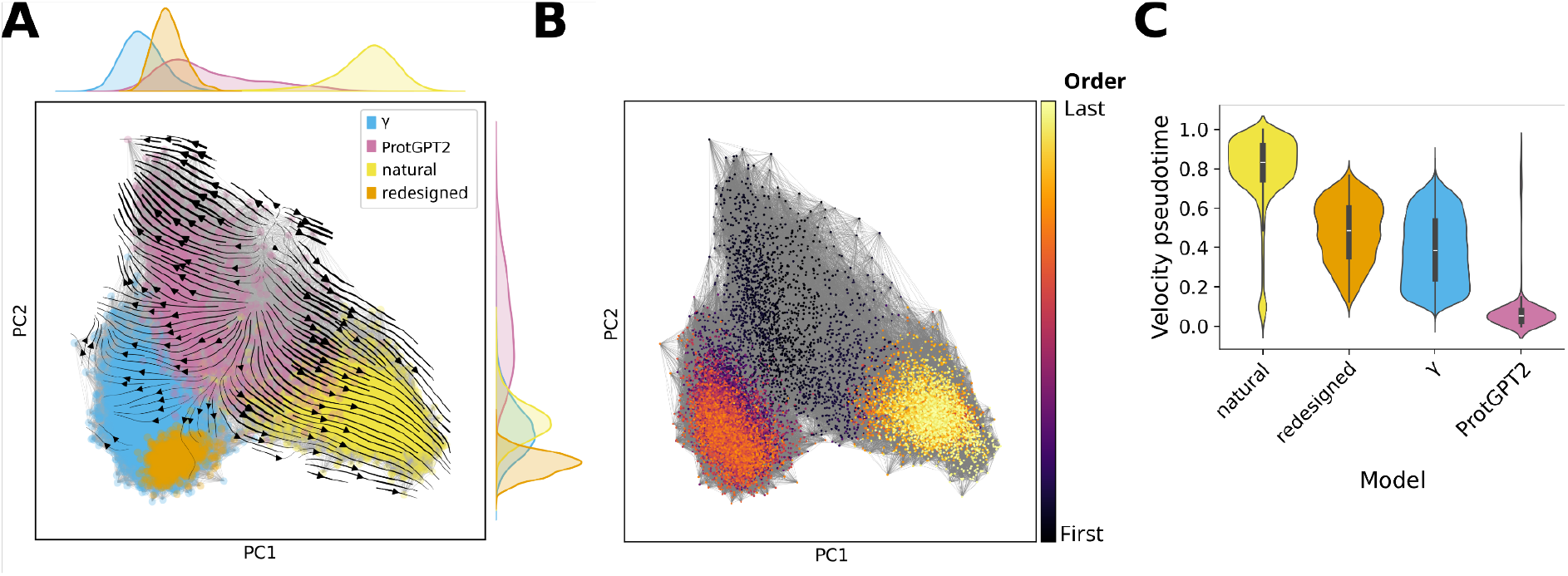
Embedding and evolutionary trajectories of zinc-finger sequences. All sequences similar to the reference zinc-finger structure were embedded using ESM-1b. Each point represents a sequence. (A) Sequences colored by type: ProtGPT2 (pink), M-ProtGPT2-γ (blue), ProteinMPNN-redesigned (orange), and natural sequences from the corresponding MSA (yellow). Clustering reflects relative positions in embedding space. (B) Sequences colored by Evolocity pseudotime, with darker colors (purple/black) indicating earlier positions and lighter colors (yellow) indicating later positions along the inferred evolutionary trajectories. (C) Violin plots showing the distribution of pseudotime for each sequence type: natural sequences (yellow), ProteinMPNN-redesigned (orange), M-ProtGPT2-γ (blue), and ProtGPT2 (pink).

Evolutionary progression along these trajectories was further characterized using velocity pseudotime, which quantifies relative position along the inferred flow from the predicted root. Pseudotime increases away from the ProtGPT2 source, with M-ProtGPT2-γ sequences occupying intermediate values between baseline and downstream states. Natural and redesigned sequences appear at later pseudotime positions, consistent with increased sequence maturity within the local embedding structure. Together, these results place γ-generated sequences between unconstrained baseline generations and sequences shaped more strongly by natural evolution or structure-guided redesign; pseudotime may also reflect increasing metal-binding specialization rather than evolutionary time alone.

To evaluate whether the regex-based motif filter was overly restrictive, we relaxed motif definitions for one representative structure per metalloprotein class (Supplementary Table S3). Spacer lengths ≥10 residues were allowed to vary by ±3 residues, and newly recovered sequences were evaluated with mymetal^34^ to estimate metal-binding probability. Only a small number of additional sequences were recovered under the relaxed definitions (Supplementary Table S4), indicating that strict regex definitions did not exclude a large pool of plausible variants.

### Exploring generated sequences beyond trivial metal-binding motifs

Sequences that do not match canonical binding motifs may still encode non-canonical or yet to discover metal-coordination patterns, representing an opportunity to explore regions of sequence space not captured by training-related motif definitions. To identify candidates with a higher likelihood of metal-binding, we implemented a multi-step filtering pipeline (Figure 5A) that progressively enriches for sequences with both high predicted structural confidence (higher pLDDT) and metal-binding potential.

**Figure 5.**
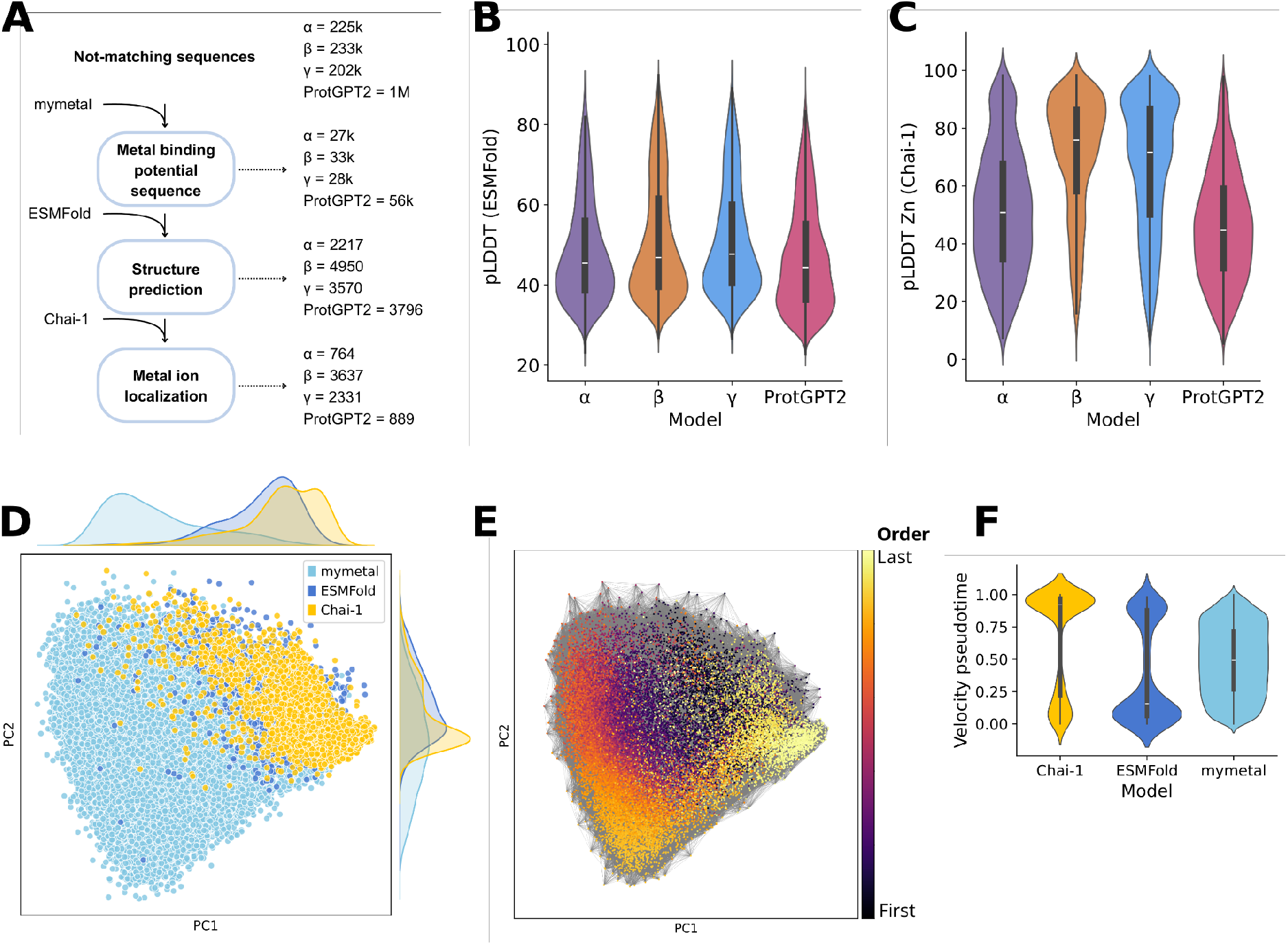
Analysis of γ-generated sequences beyond canonical motifs. (A) Schematic of the multi-step filtering pipeline applied to non–motif-matching sequences to enrich for candidates with higher predicted metal-binding likelihood. (B) Violin plots of average backbone pLDDT scores predicted by ESMFold for sequences classified as metal-binding by mymetal. (C) Violin plots of metal-ion–specific pLDDT scores predicted by Chai-1 for the subset of sequences with ESMFold average pLDDT > 70. Panels B and C compare the three M-ProtGPT2 variants (α, β, γ) with baseline ProtGPT2. (D–E) Projection of γ-generated sequences in the ESM-1b embedding space onto the first two principal components. Each point represents a sequence and is colored by (D) pipeline filtering stage or (E) Evolocity pseudotime. (F) Violin plots of pseudotime distributions for γ-generated sequences at each filtering stage, illustrating progression toward metal-binding specialization.

First, sequences were scored with mymetal^34^, retaining only those with a predicted metal-binding probability > 0.5. This step selected ~27k, 33k, and 28k sequences from the α-, β-, and γ-generated sets, representing 12–14% of the respective non-motif-matching subsets, compared to 56k (~5%) from ProtGPT2As expected, enrichment was lower than for motif-matching sequences (Supplementary Table S2), reflecting the presence of many non-metalloproteins in this set. ESMFold analysis showed that β- and γ-generated sequences had higher structural confidence than ProtGPT2 or α sequences (Figure 5B). Applying a mean pLDDT threshold of 70 retained 2,217 (α), 4,950 (β), 3,570 (γ), and 3,796 (ProtGPT2) sequences.

Metal-ion coordination was then evaluated with Chai-1. Backbone confidence was broadly comparable across models, but β- and γ-generated sequences shifted toward higher metal-ion-specific pLDDT values (Figure 5B,C). Applying a Chai-1 threshold of 60 retained 724 (α), 3,637 (β), 2,331 (γ), and 889 (ProtGPT2) sequences (Figure 5C). These results suggest that fine-tuning with larger and more chemically diverse synthetic datasets selectively enhances predicted metal-binding quality, even among sequences that do not match canonical motifs, while overall backbone folding remains broadly comparable.

To examine how the filtering pipeline shapes γ-generated sequences in embedding space, all sequences were embedded with ESM-1b and projected onto the first two principal components (Figure 5D). Sequences passing the mymetal filter were broadly distributed, whereas sequences retained by the ESMFold pLDDT threshold shifted toward higher values along PC1, reflecting enrichment for backbone structural confidence. Sequences further filtered with Chai-1 for predicted metal-binding occupy a more compact region characterized by high PC1 and central PC2 values, indicating that metal-coordination features refine the selection along an orthogonal axis (PC2). These observations suggest that PC1 primarily captures fold-related properties, while metal-binding features introduce additional structure in the embedding space.

Directional relationships inferred with Evolocity reveal complementary trends (Figure 5E,F). Pseudotime for ESMFold-selected sequences exhibits a bimodal distribution. Chai-1-selected sequences remained bimodal but showed a marked shift of the dominant peak toward higher pseudotime values, indicating progressive enrichment for metal-binding specialization along the inferred trajectory. Together, these results indicate that predicted structural confidence and metal-binding potential are partially decoupled: folding confidence organizes sequences along PC1, while metal-coordination properties are reflected in pseudotime, revealing an orthogonal dimension of functional specialization within the γ-generated subset.

### Identification of non-trivial metal-binding motifs in four-helical bundles

To evaluate whether the pipeline can uncover metal-binding sites beyond trivial motifs, we focused on four-helical bundles (4hb) as a stringent test case. This scaffold is prevalent in both natural metalloproteins^48–51^ (e.g. EXXH or HXXX[E,H] motifs) and de novo designs^52,53^, making it challenging to detect genuinely novel coordination sites.

Candidate 4hb sequences were identified among non-motif-matching sequences that passed the Chai-1 metal-ion confidence filter. Predicted structures were clustered with Foldseek, centroids were ranked by α-helical content, and top centroids were visually screened for 4hb topology. This yielded six candidate bundles for detailed analysis (Figure 6A)

**Figure 6.**
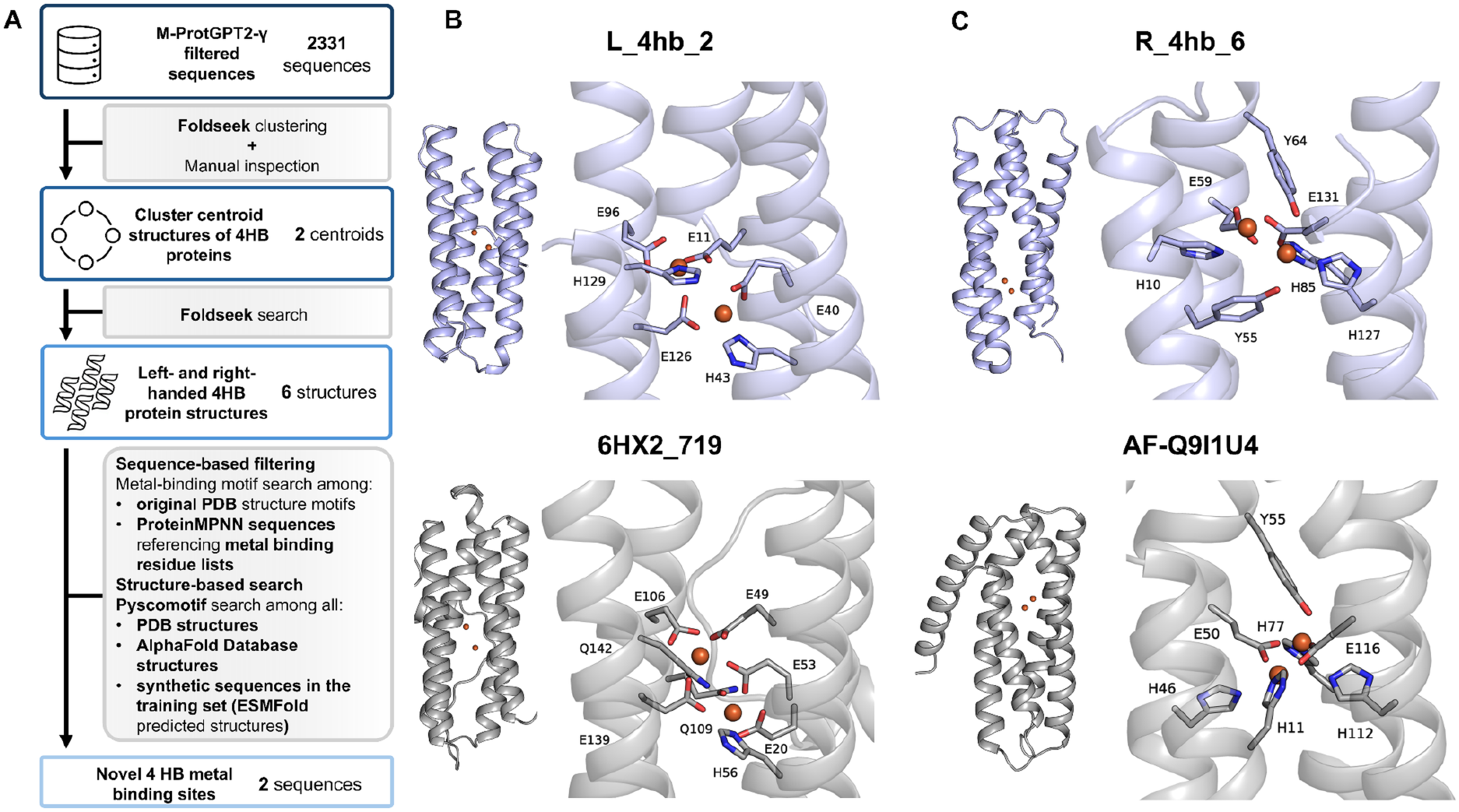
Validation of novel four-helical bundle metal-binding sites. (A) Schematic of the multi-level novelty assessment applied to four-helical bundle sequences from non–motif-matching sequences. (B) Cartoon representation of the minimized structures of di-iron proteins L_4hb_2 (top) and its closest match 6HX2_719 (bottom) and (C) R_4hb_6 (top) and its closest match AF-Q9I1U4 (bottom), with closeups of the respective metal sites. Iron(II) ions are represented as orange spheres and metal binding residues as sticks.

The identified bundles fell into two topological classes: left-handed (ferritin-like) and right-handed (hemerythrin-like). Other architectures, such as ROP-like folds, bisecting bundles, or parallel arrangements, were not recovered, possibly due to underrepresentation in the training set or centroid selection bias. Thus, the analysis does not exclude the potential existence of additional bundle types in the generated sequences.

Metal-binding motifs were extracted from Chai-1-predicted structures, modeled with either a dinuclear Fe(III) site or mononuclear Zn(II) ion. We rationalized that both mononuclear and dinuclear metal sites can be found in nature, as seen for the hemerythrin-like domain^48^. Therefore, all metal sites were subjected to a multi-level novelty assessment (Figure 6A), which also led to mononuclear/dinuclear assignment based on structural similarity. We compared them both against motifs in the original PDB training set and in the ProteinMPNN-designed training sequences using only Chai-1-assigned coordinating positions (Supplementary Tables S5-6). Then, we performed structure-based searches with Pyscomotif^54^ against the full PDB, the AlphaFold Database (AFDB)^55^, and ESMFold-predicted training-derived sequences (Supplementary Tables S7–9)

Among the six 4hb structures analyzed, two contained coordination motifs with no exact match in the PDB, AFDB, or training data. These were: (i) sequence L_4hb_2, a left-handed ferritin-like bundle with motif E11–E96–E126–H129, and (ii) sequence R_4hb_6, a right-handed hemerythrin-like bundle with motif H10–Y55–E59–H127– E131. Both were consistent with dinuclear clusters, according to RMSD, and their closest Pyscomotif matches differed by at least one coordinating residue, often replaced by non-coordinating side chains (Val, Leu, Gln).

We performed energy minimization of both L_4hb_2 and R_4hb_6 in explicit solvent and confirmed that the predicted coordination geometry is compatible with physically reasonable sites. For L_4hb_2, the closest match in the training set is sequence 6HX2_719, derived from the ferritin-like protein Dps^56^ (Figure 6B). Notably, the diiron metal-binding site in 6HX2_719 was not present in the original PDB structure but was instead introduced within the bundle core by ProteinMPNN, which hallucinated the coordination environment typical of ferritins during sequence inference. The site contains two coordinating glutamine residues (Q109, Q142), rather than the single glutamine found in natural ferritins, along with five glutamate residues and one histidine residue. While this represents the best structural match identified by Pyscomotif based on Cα positions of the metal-binding motif in L_4hb_2, the coordination composition more closely resembles that of bacterioferritin, methane monooxygenase, Δ^9^-desaturase and related proteins, which typically feature two histidine residues and four glutamate residues coordinating the diiron center. For R_4hb_6, the closest AFDB match (AF-Q9I1U4) exhibits a His→Tyr substitution at a coordinating position, preserving overall coordination but altering the chemical character of the ligand (Figure 6C). This AFDB entry corresponds to a hypothetical hemerythrin-like protein with unknown function from *P. aeruginosa*, making any further consideration purely speculative. Nevertheless, the closest structural match in the ProteinMPNN training set is 6TYJ_259 (Supplementary Figure S4), the latter being a dinuclear-binding hemerythrin from *M. kansasii* still featuring a Tyr hydroxyl moiety at 3.2 Å from one of the metals.

## 3. Discussion

A central question motivating this work is whether structure-derived synthetic data can guide protein language models (pLMs) toward specialized functional classes without compromising the broad sequence knowledge acquired during pretraining. Fine-tuning ProtGPT2 on synthetic metalloproteins shifted generation toward a distinct region of embedding space that partially overlapped ProteinMPNN redesigns while remaining separate from baseline ProtGPT2 and natural sequences.

Embedding and Evolocity analyses of sequences containing known metal-binding motifs reveal that the observed shift in generative behavior is structured rather than diffuse. Evolocity flows originating from baseline ProtGPT2 bifurcate toward natural sequences on one hand and toward synthetic redesigns and γ-generated sequences on the other. Pseudotime ordering along these trajectories does not represent biological evolutionary time but instead reflects progression along directions of increasing model-assigned likelihood within a sequence-derived embedding space. Since these embeddings are learned from sequence data alone, the inferred ordering captures intrinsic constraints encoded in protein sequences rather than explicit evolutionary history^57^, or their functional fitness^58^. In this setting, the strong association between pseudotime and enrichment for metal-binding features suggests that pseudotime primarily reflects progression in metal-binding likelihood, with γ-generated sequences occupying an intermediate position along this axis. Together, these patterns indicate that synthetic fine-tuning integrates structural and functional constraints from the training data while preserving diversity, producing a specialized branch of sequence space without erasing pretrained knowledge. This behavior parallels observations from representation-shift studies in language models, where supervised fine-tuning reorient latent features while largely preserving pretrained geometry^59,60^.

Sequences outside canonical motifs showed a similar hierarchy. Well-folded sequences occupied a coherent region of embedding space, and candidates with confident metal-binding sites formed a narrower subspace within it. This nesting suggests that metal-binding specialization is layered onto structural competence rather than learned independently, consistent with the idea that metal-binding sites often arise by refinement of pre-existing ^61–63^.

From a mechanistic perspective, synthetic data generated by inverse-folding models can be viewed as a mechanism for injecting targeted structural priors into sequence-only language models. This strategy parallels recent approaches combining structure-based generative models with protein language models for binding-pocket design, which similarly demonstrate that structural constraints integrated into sequence generation yield higher-quality functional sites than either modality alone^64^. Rather than replacing regularities learned from natural sequences, fine-tuning biases sampling toward specialized regions of sequence space while maintaining flexibility. The zinc-finger analysis illustrates this point: synthetic sequences were optimized for metal coordination but were not constrained by the natural DNA-binding functions of the family, so their trajectory remained distinct from natural zinc-finger proteins. In this framework, γ-generated sequences occupy an intermediate position between baseline and redesign distributions.

The four-helical bundle case study further shows that the model can explore underrepresented sequence space without losing fold integrity. Two of six high-confidence candidates carried coordination motifs absent from the training set, PDB, and AFDB, supporting the potential of synthetic fine-tuning for exploratory metalloprotein design.

### Limitations and outlook

Several limitations should be acknowledged. First, all analyses are computational, experimental validation is required. Second, the multi-step filtering pipeline may discard valid sequences with non-canonical binding sites, and predictive tools such as mymetal, ESMFold, and Chai-1 have intrinsic uncertainties. Third, fine-tuning was performed using a standard full-model approach. Alternative strategies, including parameter-efficient fine-tuning^13^, adapters^65,66^, reinforcement learning^67^, or Direct Preference Optimization^68^, could better preserve pretrained knowledge while improving specialization. Fourth, although ProteinMPNN is effective at generating sequences compatible with a given fold, it is not explicitly optimized to preserve functional properties, raising the possibility that synthetic sequences may emphasize structural compatibility over functional fidelity. Assessing how such biases interact with, or potentially reshape, functional representations in the fine-tuned pLM will require further investigation. Finally, the study focused on metalloproteins, in particular zinc fingers and four-helical bundles; the generalizability of these findings to other folds or functional classes remains to be evaluated.

Despite these caveats, our results show that synthetic data can guide pLMs toward targeted protein classes while preserving diversity. Extending this framework to additional metals, folds, and functions may help map underrepresented regions of protein sequence space and clarify how pLMs encode structure-function relationships.

## 4. MATERIALS AND METHODS

### 4.1 Synthetic data generation

Crystal structures were retrieved from the Protein Data Bank (PDB)^69^. The PDB advanced search interface was used to identify structures containing transition metals as chemical components (Co, Ni, Fe, Mn, Cu, Zn, and Cd).

The α dataset was constructed from approximately 370 monomeric protein structures shorter than 200 residues and containing at least one metal-binding site. A metal-binding site was defined as a metal ion coordinated by at least three side-chain heteroatoms from Cys, Asp, Glu, His, or Met residues within 2.8 Å, contributed by a single polypeptide chain.

The β dataset was assembled using the same structural and coordination criteria, with the inclusion of alkaline earth metals (Ca and Mg) in the initial PDB search, resulting in approximately 2,000 structures.

The γ dataset was generated from the same structural set as β but with substantially increased sequence sampling per structure in order to expand sequence diversity while preserving backbone constraints.

Structures containing missing backbone segments or metal-binding sites formed by residues from multiple chains were excluded from all datasets.

Fixed-backbone sequence design was performed using ProteinMPNN (soluble model weights) with a sampling temperature of 0.2. Residues directly involved in metal coordination (defined as having any side-chain heteroatom within 2.8 Å of a metal ion) were held fixed during design. For the α dataset, 200 sequences were generated per structure; for the β dataset, 100 sequences per structure; and for the γ dataset, 1,000 sequences per structure.

Sequences generated from each structural set were randomly shuffled to create the final datasets, comprising approximately 70,000 sequences for α, 140,000 sequences for β, and 1,400,000 sequences for γ.

### 4.2 Model set up and training

The three synthetic datasets were used independently to fine-tune ProtGPT2^1^, yielding three specialized variants denoted M-ProtGPT2-α, M-ProtGPT2-β, and M-ProtGPT2-γ. All model parameters were unfrozen and trained for 1–2 epochs at low learning rate (1×10^−5^ to 1×10^−6^) with HuggingFaceTransformers^70^ library using the official ProtGPT2 implementation ([github link](https://github.com/huggingface/transformers/blob/main/examples/pytorch/language-modeling/run_clm.py)). Sequences were tokenized with the original ProtGPT2 tokenizer.

Training/validation splits were performed at the PDB-structure level: 10% of scaffolds were reserved for validation and 90% for training, preventing related sequences from the same template from inflating validation performance.

Fine-tuning was carried out on four NVIDIA Tesla V100 GPUs (32 GB memory each) with a batch size of 32. Multi-GPU training was orchestrated using DeepSpeed in combination with the Accelerate library.

For generation, fine-tuned weights were loaded into the HuggingFace text-generation pipeline with maximum length 70, repetition penalty 1.5, do_sample = True, and top_k = 950. Approximately 1.5 million sequences were generated per model in batches of 200 with seed resets between batches. Invalid amino acid tokens were removed during post-processing.

For baseline comparison, approximately 1.5 million sequences were generated using the original ProtGPT2 model under identical sampling parameters and filtering criteria.

### 4.3 Generated sequences analysis

#### Motif-based sequence classification and structural analysis of motif-matching sequences

Metal-binding motifs present in the training set were encoded as regular expressions (regex). Each motif specifies coordinating residues (e.g., Cys, His, Glu) separated by variable-length spacers corresponding to the observed residue intervals. Generated sequences were screened against these regex patterns and classified as motif-matching or non-motif-matching.

Motif-matching analyses focused on γ-generated sequences, with motif-matching baseline ProtGPT2 sequences processed identically for comparison. Only motifs derived from selected reference proteins were retained, spanning iron-sulfur proteins, zinc fingers, blue copper proteins, oxidases, and hydrolases (Supplementary Table S1).

Filtered sequences were subjected to structure prediction using ESMFold. Mean pLDDT scores were computed for each predicted structure. Structural similarity was evaluated using Foldseek (easy-search --max-seqs 10000 -s 10), with each reference PDB structure as query and predicted structures as targets. Sequence similarity was assessed using MMseqs2^71^ (easy-search -s 7.0 --cov-mode 0 --alignment-mode 3), and TM-scores were compared to sequence identity (fident).

Evolocity was applied to γ-generated sequences, ProtGPT2 baseline sequences, ProteinMPNN redesigns, and natural sequences. For the zinc-finger class, sequences with TM-scores > 0.5 relative to their corresponding reference structures were retained. Multiple sequence alignments matching the relevant motifs were retrieved. When necessary, natural sequences were randomly subsampled to match the number of generated sequences prior to Evolocity analysis.

#### Multi-step filtering pipeline for non-motif-matching sequences

Non-motif-matching sequences were filtered sequentially by mymetal^34^ score (> 0.5), ESMFold mean pLDDT (> 70), and Chai-1 metal-ion-specific pLDDT (> 60) using a single Zn2+ ion model.

Evolocity analysis was performed on γ-generated sequences at three stages of the filtering pipeline: (i) sequences predicted as metal-binding by mymetal (> 0.5), (ii) sequences with ESMFold mean pLDDT > 70, and (iii) sequences with Chai-1 metal ion pLDDT > 60. For each subset, evolutionary vector fields and pseudotime distributions were computed.

To test the impact of strict regex definitions, relaxed motifs were built for one representative PDB structure per metalloprotein class. For motifs with spacer regions ≥10 residues, spacer lengths were expanded by ±3 residues, and sequences recovered only under these relaxed rules were evaluated with mymetal.

#### Four-helical bundle identification and characterization

Pipeline-filtered structures were clustered with Foldseek (-c 0.9), centroids were ranked by helical content, and top clusters were visually screened for four-helical bundle topologies. Representative bundles were then used in additional Foldseek searches to recover related architectures. Metal-binding motifs extracted from Chai-1-predicted 4hb structures modeled with either two Fe(III) ions or one Zn(II) ion were searched against PDB training structures and synthetic training sequences. Matches were counted only when aligned residues corresponded to annotated metal-binding positions in the original PDB structures (Supplementary Tables S5-S6).

Structure-based searches using Pyscomotif^54^ were conducted against the full PDB (Table S7) and AlphaFold Database (Table S8) using pre-built indexes. To exclude pre-existing motifs in the training set, four-helix bundle proteins were identified using Foldseek searches against reference structures: right-handed bundle (hemerythrin, PDB 4XPX) and left-handed bundle (ferritin, PDB 8IQV). Structures with E-value <0.2 and query coverage >70% were selected. All synthetic sequences from matching structures (73,000 sequences) were collected, predicted with ESMFold v1 (4 recycles), and indexed for custom Pyscomotif searches (Table S9).

Finally, 5 solvated systems were prepared for energy minimization: (i) γ-generated 4hb structures L_4hb_2, R_4hb_6, and (ii) their closest ProteinMPNN(6HX2_719, 6TYJ_259) or AFDB match (AF-Q9I1U4), both with two Fe^2+^ atoms in the metal-binding site. Structures were solvated using CHARMM-GUI in octahedral boxes with 10 Å buffer, neutralized with K^+^or Cl^−^ ions, and minimized using NAMD3 for 30,000 conjugated gradient steps with applied constraints.

## Data and code availability

All original code and data used in our analysis has been deposited to Zenodo and are publicly available as of the date of publication (https://zenodo.org/records/18672158; https://doi.org/10.5281/zenodo.18672158). Our fine-tuned models are available on HuggingFace at https://huggingface.co/gsgueglia. Any additional information required to reanalyze the data reported in this paper is available from the lead contact upon request.

## Declaration of Generative AI and AI-assisted technologies

During the preparation of this manuscript, the authors employed generative AI tools, including ChatGPT (OpenAI), Claude (Anthropic) and Gemini (Google), for language refinement. These tools were used solely to improve clarity, coherence, and style. All AI-assisted content was carefully reviewed and substantively edited by the authors, who accept full responsibility for the accuracy and integrity of the final submitted work.

## References

1. Ferruz, N., Schmidt, S. & Höcker, B. ProtGPT2 is a deep unsupervised language model for protein design. Nat. Commun. 13, 4348 (2022).

2. Nijkamp, E., Ruffolo, J. A., Weinstein, E. N., Naik, N. & Madani, A. ProGen2: Exploring the boundaries of protein language models. Cell Syst. 14, 968–978.e3 (2023).

3. Lin, Z. et al. Evolutionary-scale prediction of atomic-level protein structure with a language model. Science 379, 1123–1130 (2023).

4. Ruffolo, J. A. & Madani, A. Designing proteins with language models. Nat. Biotechnol. 42, 200–202 (2024).

5. Bepler, T. & Berger, B. Learning the protein language: Evolution, structure, and function. Cell Syst. 12, 654–669.e3 (2021).

6. Madani, A. et al. Large language models generate functional protein sequences across diverse families. Nat. Biotechnol. 41, 1099–1106 (2023).

7. Jumper, J. et al. Highly accurate protein structure prediction with AlphaFold. Nature 596, 583–589 (2021).

8. Wu, R. et al. High-resolution de novo structure prediction from primary sequence. 2022.07.21.500999 Preprint at 10.1101/2022.07.21.500999 (2022).

9. Discovery, C. et al. Chai-1: Decoding the molecular interactions of life. 2024.10.10.615955 Preprint at 10.1101/2024.10.10.615955 (2024).

10. Li, X., Perez, R., Ferrie, J. J., Petersson, E. J. & Giannakoulias, S. Accurate Prediction of Protein Tertiary and Quaternary Stability Using Fine-Tuned Protein Language Models and Free Energy Perturbation. Int. J. Mol. Sci. 26, 7125 (2025).

11. Rekapalli, B., Wuichet, K., Peterson, G. D. & Zhulin, I. B. Dynamics of domain coverage of the protein sequence universe. BMC Genomics 13, 634 (2012).

12. Unger, R., Uliel, S. & Havlin, S. Scaling law in sizes of protein sequence families: from super-families to orphan genes. Proteins 51, 569–576 (2003).

13. Sledzieski, S. et al. Democratizing protein language models with parameter-efficient fine-tuning. Proc. Natl. Acad. Sci. 121, e2405840121 (2024).

14. Gholami, S. & Omar, M. Does Synthetic Data Make Large Language Models More Efficient? Preprint at 10.48550/arXiv.2310.07830 (2023).

15. Li, H. et al. Synthetic Data (Almost) from Scratch: Generalized Instruction Tuning for Language Models. Preprint at 10.48550/arXiv.2402.13064 (2024).

16. Liu, R. et al. Best Practices and Lessons Learned on Synthetic Data. Preprint at 10.48550/arXiv.2404.07503 (2024).

17. Seddik, M. E. A., Chen, S.-W., Hayou, S., Youssef, P. & Debbah, M. How Bad is Training on Synthetic Data? A Statistical Analysis of Language Model Collapse. Preprint at 10.48550/arXiv.2404.05090 (2024).

18. Wang, Z. et al. CodecLM: Aligning Language Models with Tailored Synthetic Data. Preprint at 10.48550/arXiv.2404.05875 (2024).

19. Mormille, L. H., Salama, I. & Atsumi, M. Domain-Specific Vision Transformer Pre-Training with Synthetic Data. in Proceedings of the 2024 8th International Conference on Advances in Artificial Intelligence 322–326 (Association for Computing Machinery, New York, NY, USA, 2025). doi:10.1145/3704137.3704168.

20. Sampath, V., Maurtua, I., Aguilar Martín, J. J. & Gutierrez, A. A survey on generative adversarial networks for imbalance problems in computer vision tasks. J. Big Data 8, 27 (2021).

21. Dauparas, J. et al. Robust deep learning–based protein sequence design using ProteinMPNN. Science 378, 49–56 (2022).

22. Krapp, L. F. et al. Context-aware geometric deep learning for protein sequence design. Nat. Commun. 15, 6273 (2024).

23. Holm, R. H., Kennepohl, P. & Solomon, E. I. Structural and Functional Aspects of Metal Sites in Biology. Chem. Rev. 96, 2239–2314 (1996).

24. Andreini, C., Bertini, I., Cavallaro, G., Holliday, G. L. & Thornton, J. M. Metal ions in biological catalysis: from enzyme databases to general principles. JBIC J. Biol. Inorg. Chem. 13, 1205–1218 (2008).

25. Li, J. et al. The Metal-binding Protein Atlas (MbPA): An Integrated Database for Curating Metalloproteins in All Aspects. J. Mol. Biol. 435, 168117 (2023).

26. Putignano, V., Rosato, A., Banci, L. & Andreini, C. MetalPDB in 2018: a database of metal sites in biological macromolecular structures. Nucleic Acids Res. 46, D459–D464 (2018).

27. Jeong, W. J., Lee, J., Eom, H. & Song, W. J. A Specific Guide for Metalloenzyme Designers: Introduction and Evolution of Metal-Coordination Spheres Embedded in Protein Environments. Acc. Chem. Res. 56, 2416–2425 (2023).

28. Mann, S. I., Heinisch, T., Ward, T. R. & Borovik, A. S. Coordination chemistry within a protein host: regulation of the secondary coordination sphere. Chem. Commun. Camb. Engl. 54, 4413–4416 (2018).

29. Shook, R. L. & Borovik, A. S. Role of the Secondary Coordination Sphere in Metal-Mediated Dioxygen Activation. Inorg. Chem. 49, 3646–3660 (2010).

30. Lin, G.-Y., Su, Y.-C., Huang, Y. L. & Hsin, K.-Y. MESPEUS: a database of metal coordination groups in proteins. Nucleic Acids Res. 52, D483–D493 (2024).

31. Yang, J., Roy, A. & Zhang, Y. BioLiP: a semi-manually curated database for biologically relevant ligand–protein interactions. Nucleic Acids Res. 41, D1096–D1103 (2013).

32. Haberal, i. & Oğul, H. Prediction of Protein Metal Binding Sites Using Deep Neural Networks. Mol. Inform. 38, 1800169 (2019).

33. Sánchez-Aparicio, J.-E. et al. BioMetAll: Identifying Metal-Binding Sites in Proteins from Backbone Preorganization. J. Chem. Inf. Model. 61, 311–323 (2021).

34. Aptekmann, A. A. et al. mebipred: identifying metal-binding potential in protein sequence. Bioinformatics 38, 3532–3540 (2022).

35. Lu, C.-H. et al. MIB2: metal ion-binding site prediction and modeling server. Bioinformatics 38, 4428–4429 (2022).

36. Krapp, L. F., Abriata, L. A., Cortés Rodriguez, F. & Dal Peraro, M. PeSTo: parameter-free geometric deep learning for accurate prediction of protein binding interfaces. Nat. Commun. 14, 2175 (2023).

37. Dürr, S. L., Levy, A. & Rothlisberger, U. Metal3D: a general deep learning framework for accurate metal ion location prediction in proteins. Nat. Commun. 14, 2713 (2023).

38. Zheng, H. et al. PinMyMetal: A hybrid learning system to accurately model metal binding sites in macromolecules. Res. Sq. rs.3.rs-3908734 (2024) doi:10.21203/rs.3.rs3908734/v1.

39. Shenoy, A., Kalakoti, Y., Sundar, D. & Elofsson, A. M-Ionic: prediction of metal-ion-binding sites from sequence using residue embeddings. Bioinformatics 40, btad782 (2024).

40. Lin, X. et al. SuperMetal: A Generative AI Framework for Rapid and Precise Metal Ion Location Prediction in Proteins. bioRxiv 2025.03.21.644685 (2025) doi:10.1101/2025.03.21.644685.

41. Dürr, S. L. & Rothlisberger, U. AllMetal3D: joint prediction of localization, identity and coordination geometry of common metal ions in proteins. 2025.02.05.636627 Preprint at 10.1101/2025.02.05.636627 (2025).

42. Sgueglia, G., Vrettas, M. D., Chino, M., De Simone, A. & Lombardi, A. MetalHawk: Enhanced Classification of Metal Coordination Geometries by Artificial Neural Networks. J. Chem. Inf. Model. 64, 2356–2367 (2024).

43. Hekkelman, M. L., de Vries, I., Joosten, R. P. & Perrakis, A. AlphaFill: enriching AlphaFold models with ligands and cofactors. Nat. Methods 20, 205–213 (2023).

44. Dai, X., Henderson, M., Yoo, S. & Liu, Q. Predicting Metal-binding Proteins and Structures Through Integration of Evolutionary-scale and Physics-based Modeling. J. Mol. Biol. 437, 168962 (2025).

45. Feehan, R., Franklin, M. W. & Slusky, J. S. G. Machine learning differentiates enzymatic and non-enzymatic metals in proteins. Nat. Commun. 12, 3712 (2021).

46. van Kempen, M. et al. Fast and accurate protein structure search with Foldseek. Nat. Biotechnol. 1–4 (2023) doi:10.1038/s41587-023-01773-0.

47. Hie, B. L., Yang, K. K. & Kim, P. S. Evolutionary velocity with protein language models predicts evolutionary dynamics of diverse proteins. Cell Syst. 13, 274–285.e6 (2022).

48. Alvarez-Carreño, C., Alva, V., Becerra, A. & Lazcano, A. Structure, function and evolution of the hemerythrin-like domain superfamily. Protein Sci. 27, 848–860 (2018).

49. Pozzi, C. et al. Iron binding to human heavy-chain ferritin. Acta Crystallogr. D Biol. Crystallogr. 71, 1909–1920 (2015).

50. Pope, S. R. et al. Heme Oxygenase–Like Metalloenzymes. Annu. Rev. Biochem. 94, 59–88 (2025).

51. Yoon, J., Fujii, S. & Solomon, E. I. Geometric and electronic structure differences between the type 3 copper sites of the multicopper oxidases and hemocyanin/tyrosinase. Proc. Natl. Acad. Sci. 106, 6585–6590 (2009).

52. Chalkley, M. J., Mann, S. I. & DeGrado, W. F. De novo metalloprotein design. Nat. Rev. Chem. 6, 31–50 (2022).

53. Lombardi, A., Pirro, F., Maglio, O., Chino, M. & DeGrado, W. F. De Novo Design of Four-Helix Bundle Metalloproteins: One Scaffold, Diverse Reactivities. Acc. Chem. Res. 52, 1148–1159 (2019).

54. Cia, G., Kwasigroch, J., Stamatopoulos, B., Rooman, M. & Pucci, F. pyScoMotif: discovery of similar 3D structural motifs across proteins. Bioinforma. Adv. 3, vbad158 (2023).

55. Fleming, J. et al. AlphaFold Protein Structure Database and 3D-Beacons: New Data and Capabilities. J. Mol. Biol. 437, 168967 (2025).

56. Zeth, K., Sancho-Vaello, E. & Okuda, M. Metal Positions and Translocation Pathways of the Dodecameric Ferritin-like Protein Dps. Inorg. Chem. 58, 11351–11363 (2019).

57. Hie, B., Zhong, E. D., Berger, B. & Bryson, B. Learning the language of viral evolution and escape. Science 371, 284–288 (2021).

58. Luo, Y. et al. ECNet is an evolutionary context-integrated deep learning framework for protein engineering. Nat. Commun. 12, 5743 (2021).

59. Khayatan, P., Shukor, M., Parekh, J., Dapogny, A. & Cord, M. Analyzing Finetuning Representation Shift for Multimodal LLMs Steering. Preprint at 10.48550/arXiv.2501.03012 (2025).

60. Zhou, Y. & Srikumar, V. A Closer Look at How Fine-tuning Changes BERT. in Proceedings of the 60th Annual Meeting of the Association for Computational Linguistics (Volume 1: Long Papers) (eds Muresan, S., Nakov, P. & Villavicencio, A.) 1046–1061 (Association for Computational Linguistics, Dublin, Ireland, 2022). doi:10.18653/v1/2022.acl-long.75.

61. Bromberg, Y. et al. Quantifying structural relationships of metal-binding sites suggests origins of biological electron transfer. Sci. Adv. 8, eabj3984 (2022).

62. Dupont, C. L., Butcher, A., Valas, R. E., Bourne, P. E. & Caetano-Anollés, G. History of biological metal utilization inferred through phylogenomic analysis of protein structures. Proc. Natl. Acad. Sci. 107, 10567–10572 (2010).

63. Sendra, K. M. et al. An ancient metalloenzyme evolves through metal preference modulation. Nat. Ecol. Evol. 7, 732–744 (2023).

64. Zhang, Z., Shen, W. X., Liu, Q. & Zitnik, M. Efficient generation of protein pockets with PocketGen. Nat. Mach. Intell. 6, 1382–1395 (2024).

65. Houlsby, N. et al. Parameter-efficient transfer learning for NLP. in International Conference on Machine Learning 2790–2799 (PMLR, 2019).

66. Hu, E. J. et al. LoRA: Low-Rank Adaptation of Large Language Models. Preprint at 10.48550/arXiv.2106.09685 (2021).

67. Ouyang, L. et al. Training language models to follow instructions with human feedback. Preprint at 10.48550/arXiv.2203.02155 (2022).

68. Rafailov, R. et al. Direct Preference Optimization: Your Language Model is Secretly a Reward Model. Preprint at 10.48550/arXiv.2305.18290 (2024).

69. Berman, H. M. et al. The Protein Data Bank. Nucleic Acids Res. 28, 235–242 (2000).

70. Wolf, T. et al. HuggingFace’s Transformers: State-of-the-art Natural Language Processing. Preprint at 10.48550/arXiv.1910.03771 (2020).

71. Hauser, M., Steinegger, M. & Söding, J. MMseqs software suite for fast and deep clustering and searching of large protein sequence sets. Bioinformatics 32, 1323–1330 (2016).

